# Fast lightweight accurate xenograft sorting

**DOI:** 10.1101/2020.05.14.095604

**Authors:** Jens Zentgraf, Sven Rahmann

**Affiliations:** Bioinformatics, Computer Science XI, TU Dortmund University, 44221 Dortmund, Germany; Genome Informatics, Institute of Human Genetics, University of Duisburg-Essen, 45122 Essen, Germany

## Abstract

**Motivation:** With an increasing number of patient-derived xenograft (PDX) models being created and subsequently sequenced to study tumor heterogeneity and to guide therapy decisions, there is a similarly increasing need for methods to separate reads originating from the graft (human) tumor and reads originating from the host species’ (mouse) surrounding tissue. Two kinds of methods are in use: On the one hand, alignment-based tools require that reads are mapped and aligned (by an external mapper/aligner) to the host and graft genomes separately first; the tool itself then processes the resulting alignments and quality metrics (typically BAM files) to assign each read or read pair. On the other hand, alignment-free tools work directly on the raw read data (typically FASTQ files). Recent studies compare different approaches and tools, with varying results.

**Results:** We show that alignment-free methods for xenograft sorting are superior concerning CPU time usage and equivalent in accuracy. We improve upon the state of the art sorting by presenting a fast lightweight approach based on three-way bucketed quotiented Cuckoo hashing. Our hash table requires memory comparable to an FM index typically used for read alignment and less than other alignment-free approaches. It allows extremely fast lookups and uses less CPU time than other alignment-free methods and alignment-based methods at similar accuracy.

**Availability:** Our software *xengsort* is available under the MIT license at http://gitlab.com/genomeinformatics/xengsort. It is written in numba-compiled Python and comes with Snakemake workflows for hash table construction and dataset processing.

## 1 Introduction

To learn about tumor heterogeneity and tumor progression under realistic *in vivo* conditions, but without putting human life at risk, one can implant human tumor tissue into a mouse and study its evolution. This is called a (patient-derived) xenograft (PDX). Over time, several samples of the (graft / human) tumor and surrounding (host / mouse) tissue are taken and subjected to exome or whole genome sequencing in order to monitor the changing genomic features of the tumor. This information can be used to predict the response to different chemotherapy alternatives and to monitor treatment success or failure. A key step in such analyses is *xenograft sorting*, i.e., separating the human tumor reads from the mouse reads. A recent study (1) showed that if such a step is omitted, several mouse reads would be aligned to certain regions of the human genome (HAMA: human-aligned mouse allele) and induce false positive variant calls for the tumor; this especially concerns certain oncogenes.

Several tools have been developed for xenograft sorting, motivated by different goals and using different approaches; a summary appears below. Here we improve upon the existing approaches in several ways: By using carefully engineered *k*-mer hash tables, our approach is both *faster* and needs *less memory* than existing tools. By designing a new decision function, we also obtain *fewer unclassified reads* and in some cases even *higher classification accuracy*. Since we use a comprehensive reference of the genome and transcriptome, we are able to uniformly deal with genome, exome, and transcriptome data of xenografts.

Concerning related work, we distinguish alignmet-based methods that work on already aligned reads (BAM files), versus alignment-free methods that directly work on short subsequences (*k*-mers) of the raw reads (FASTQ files). On the other hand, we do not distinguish between the type of data that the tools have been applied to (transcripts, or genomic DNA), because this does not depend so much on the tool but rather on the reference sequences used (genome, exome, set of transcripts, etc.).

Alignment-based methods scan existing alignments in BAM files and test whether each read maps better to the graft or to the host genome. Differences result from different parameter settings used for the alignment tool (often `bwa` or `bowtie2`) and from the way “better alignment” is defined by each of these tools. Alignment-free methods use a large lookup table to associate species information with each *k*-mer.

In Table 1, we list properties of existing tools and of *xengsort*, our implementation of the method we describe in this article. These tools support different operations: Operation “count” outputs proportions of reads belonging to each category (host, graft, etc.); operation “sort” sorts reads or alignments into different files according to origin, ideally into five categories: host, graft, both, neither, ambiguous; a “partial sort” only has three categories: host, graft, both/other; operation “filter” writes only an output file with graft reads or alignments. The sort operation is more general than the filter or partial sort operation and allows full flexibility in downstream processing. When available, the count operation is faster than counting the output of the sort operation, because it avoids the overhead of creating new BAM or FASTQ files. *XenofilteR, Xenosplit, Bamcmp* and *Disambiguate* all work on aligned BAM files. This means that the reads must be mapped and aligned with a supported read mapper first (typically, ‘bwa mem’) and the resulting BAM file must be sorted in a specific way required by the tool. The tool is typically a script that reads and compares the mapping scores and qualities in the two BAM files containing host and graft alignments. In principle, all of these tools do the same thing; large differences result rather from different alignment parameters than the tool itself. We therefore picked *XenofilteR* as a representative of this family, also because it performed well in a recent comparison (1).

**Table 1.** Tools for xenograft sorting and read filtering with key properties. See text for definition of operations; Lang.: programming language.

| Tool | Ref. | Input | Operations | Lang. |
| --- | --- | --- | --- | --- |
| *XenofilteR* | (2) | aligned BAM | filter | R |
| *Xenosplit* | (3) | aligned BAM | filter, count | Python |
| *Bamcmp* | (4) | aligned BAM | partial sort | C++ |
| *Disambiguate* | (5) | aligned BAM | partial sort | Python or C++ |
| *BBsplit* | (6) | raw FASTQ | partial sort | Java |
| *xenome* | (7) | raw FASTQ | count, sort | C++ |
| *xengsort* | (this) | raw FASTQ | count, sort | Python + numba |

*BBsplit* (part of BBTools) is special in the sense that it performs the read mapping itself, against multiple references simultaneously, based on *k*-mer seeds. Unfortunately, only up to approximately 1.9 billion *k*-mers can be indexed because of Java’s array indexing limitations (up to 2^31^ elements) and a table load limit of 0.9; so BBsplit was not usable for our human-mouse index that contains approximately 4.5 · 10^9^ *>* 2^32^ *k*-mers.

The tool *xenome* (7) is similar to our approach: It is based on a large hash table of *k*-mers and sorts the reads into several categories (host, graft, both, neither, ambiguous). A read is classified based on its *k*-mer content according to relatively strict rules. We found the threading code of *xenome* to be buggy, such that the pure counting mode resulted in a deadlock and produced no output. The sorting mode produced the complete output but then did not terminate either.

Recent studies (1, 8, 9) have compared the computational efficiency of several methods, as well as the classification accuracy of these methods and the effects on subsequent variant calling after running vs. not running xenograft sorting. The results were contradictory, with some studies reporting that alignmet-based tools are more efficient than alignment-free tools, and different tools achieving highest accuracry. Our interpretation of the results of (1) is that each of the existing approaches is able to sort with good accuracy and the main difference is in computational efficiency. Results about efficiency have to be interpreted with care because sometimes the time for alignment is included and sometimes not.

## 2 Methods

### 2.1 Overview

By considering all available host and graft reference sequences (both transcripts and genomic sequences of mouse and human), we build a large key-value store that allows us to look up the species of origin (host, graft or both) of each DNA/RNA *k*-mer that occurs in either species. A sequenced dataset (a collection of single-end or paired-end FASTQ files) is then processed by iterating over reads or read pairs, looking up the species of origin of each *k*-mer in a read (host, graft, both or none) and classifying the read based on a decision rule.

Our implementation of the key-value store as a three-way bucketed Cuckoo hashtable makes *k*-mer lookup faster than in other methods; the associated value can often be retrieved with a single random memory access. A high load factor of the hash table, combined with the technique of quotienting, ensures a low memory footprint, without resorting to approximate membership data structures, such as Bloom filters.

### 2.2 Key-value stores of canonical *k*-mers

We partition the reference genome (plus alternative alleles and unplaced contigs) and transcriptome into short substrings of a given length *k* (so-called *k*-mers); we evaluated *k* ∈ {23, 25, 27}. For each *k*-mer (“key”) in any of the reference sequences, we store whether it occurs exclusively in the host referece, exclusively in the graft reference, or in both, represented by “values” 1, 2, 3, respectively. For the host- and graft-exclusive *k*-mers, we also store whether a closely similar *k*-mer (at Hamming distance 1) occurs in the other species (add value 4); such a *k*-mer is then called a *weak* (host or graft) *k*-mer. This idea extends the *k*-mer classification of *xenome* (7), where a *k*-mer can be host, graft, both, or marginal, the latter category comprising both our weak host and weak graft *k*-mers. So we store, for each *k*-mer, a value from the 5-element set “host” (1), “graft” (2), “both” (3), “weak host” (5), “weak graft” (6). This value is stored using 3 bits. While a more compact base-5 representation is possible (e.g., storing 3 values with 125 *<* 128 = 2^7^ combinations in 7 bits instead of in 9 bits), we decided to use slightly more memory for higher speed.

To be precise, we do not work on *k*-mers directly, but on their canonical integer representations (*canonical codes*), such that a *k*-mer and its reverse complement map to the same number. We use a simple base-4 numeric encoding A ↦ 0, C ↦ 1, G ↦ 2, T/U ↦ 3, e.g. reading the 4-mer AGCG as (0212)_4_ = 38 and its reverse complement CGCT as (1213)_4_ = 103. The *canonical code* is then the *maximum* of these two numbers; here the canonical code of both AGCG and CGCT is thus 103. (In *xenome*, canonical *k*-mer codes are implemented with a more complex but still deterministic function of the two base-4 encodings; in other tools, it is often the minimum of the two encodings.) For odd *k*, there are exactly *c*(*k*) := 4^*k*^*/*2 different canonical *k*-mer codes, so each can be stored in 2*k* − 1 bits in principle. However, implementing a *fast* bijection of the set of canonical codes (which is a subset of size *c*(*k*) of {0 .. (4^*k*^ − 1)}) to {0 .. (*c*(*k*) − 1)} seems difficult, so we use 2*k* bits to store the canonical code directly, which allows faster access. We do use quotienting, described below, to reduce the size; yet in principle, one additional bit could be saved.

### 2.3 Multi-way bucketed quotiented Cuckoo hashing

We use multi-way bucketed Cuckoo hash table as the data structure for the *k*-mer key-value store. Let *C* be the set of canonical codes of *k*-mers; as explained above, we take *C* ={0 .. (4^*k*^ − 1)}, even though only half of the codes are used (for odd *k*). Let *P* be the set of locations (buckets) in the hash table and *p* their number; we set *P* := {0 .. (*p* − 1)}. Eeach key can be stored at up to *h* different locations (buckets) in the table. The possible buckets for a code are computed by *h* different hash functions *f*_1_, *f*_2_, *…, f*_*h*_ : *C* → *P*. Each bucket can store up to a certain number *b* of key-value pairs. So there is space for *N* := *pb* key-value pairs in the table overall, and each pair can be stored at one of *hb* locations in *h* buckets. Together with an insertion strategy as described below, this framework is referred to as (*h, b*) Cuckoo hashing. Classical Cuckoo hashing uses *h* = 2 and *b* = 1; for this work, we use *h* = 3 and *b* = 4. A visualization is provided in Figure 1. Using several hash functions and larger buckets increases the load limit; using *h* = 3 and *b* = 4 allows a load factor of over 99.9% (10, Table 1), while classical Cuckoo hashing only allows to fill 50% of the table.

**Fig. 1.**
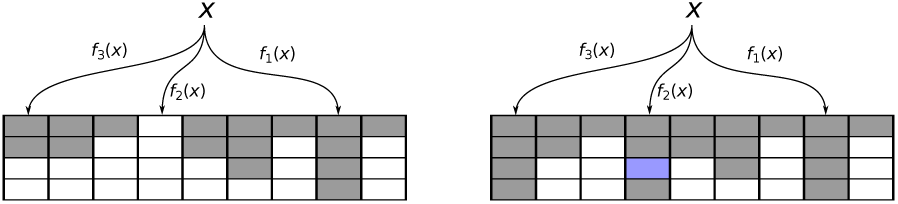
Illustration of (3,4) Cuckoo hashing with 3 hash functions and buckets of size 4. **Left:** Each key-value pair can be stored at one of up to 12 locations in 3 buckets. For key *x*, the bucket given by *f*_1_ (*x*) is full, so bucket *f*_2_ (*x*) is attempted, where a free slot is found. **Right:** If all *hb* slots are full, key *x* is placed into one of these slots at random (blue), and the currently present key-value pair is evicted and re-inserted into an alternative slot.

#### Search and insert

Searching for a key-value pair works as follows. Given key (canonical code) *x*, first *f*_1_(*x*) is computed, and this bucket is searched for key *x* and the associated value. If it is not found, buckets *f*_2_(*x*) and then *f*_3_(*x*) are searched similarly. Each bucket access is a random memory lookup and most likely triggers a cache miss. We can ensure that each bucket is contained within a single cache line (by using additional padding bits if necessary). Then, the number of cache misses is limited to *h* = 3 for one search operation.

Because we fill the table well below the load limit (at 88% of 99.9%), we are able to store most key-value pairs in the bucket indicated by the *first* hash function *f*_1_, and only incur a single cache miss when looking for them. Unsuccessful searches (for *k*-mers that are not present in either host or graft genome) will always need *h* memory accesses. However, one optimization is possible: If, say, the first bucket *f*_1_(*x*) contains an empty slot, we do not need to search further, because the insertion procedure produced a tight layout, in the sense that if a single element could be moved to an “earlier” bucket, it would have been done.

Insertion of a key-value pair works as follows. First, the key is searched as described above. If it is found, the value is updated with the new value. For example, if an exisiting host *k*-mer is to inserted again as a graft *k*-mer, the value is updated to “both”. If the key is not found, we check whether any of the buckets *f*_1_(*x*), *f*_2_(*x*), *f*_3_(*x*) contains a free slot. If this is the case, *x* and its value are inserted there. If all buckets are full, a random slot among the *hb* slots is picked, and the key-value pair stored there is evicted (like a cuckoo removes eggs from other birds’ nests) to make room for *x* and its value. Then an alternative location for the evicted element is searched. This process may continue for several iterations and is called a “random walk” through the table. If the walk becomes too long (longer than 5000 steps, say), we declare that the table is too full, and construction fails and has to be restarted with a larger table or different random seed.

We require that the final size (number of buckets *p*) of the hash table is known in advance, so we can pre-allocate it. The genome length is a good (over-)estimate of the number of distinct *k*-mers and can be used. We recently presented a practical algorithm (11) to optimize the assignment of *k*-mers to buckets (i.e., their hash function choices) such that the average search cost of present *k*-mers is minimized to the provable optimum. This optimization takes significant additional time and requires large additional data structures; so we took the opportunity here to evaluate whether it significantly improves lookup times in comparison to a table filled by the above random walk strategy.

#### Bijective hash functions and quotienting

In principle, we need to store the 2*k* bits for the canonical *k*-mer code *x* and the 3 bits for the value at each slot. However, by using hash functions of the form *f* (*x*) := *g*(*x*) mod *p*, where *p* is the number of buckets and *g* is a *bijective* (randomized) transformation on the full key set {0 .. (4^*k*^ − 1)}, we can encode part of *x* in *f* (*x*): Note that from *f* (*x*) and *q*(*x*) := *g*(*x*)*//p* (integer division), we can recover *g*(*x*) = *p· q*(*x*)+ *f* (*x*), and since *g* is bijective, we can recover *x* itself. This means that we only need to store *q*(*x*), not *x* itself in bucket *f* (*x*), which only takes ⌈2*k* − log_2_ *p*⌉ instead of 2*k* bits. However, since we have *h* alternative hash functions, we also need to store *which* hash function we used, using 2 bits for *h* = 3 (0 indicating that the slot is empty). This technique is known as *quotienting*. It gives higher savings for smaller buckets (for constant *N* = *pb*, smaller *b* means larger *p*), but on the other hand the load limit is smaller for small *b*. We find *b* = 4 to be a good compromise, allowing table loads of 99.9%.

For the bijective part *g*(*x*), we use affine functions of the form 

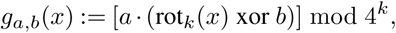

where rot_*k*_ performs a cyclic rotation of *k* bits (half the width of *x*), moving the “random” inner bits to outer positions and the less random outer bits (due to the max operation when taking canonical codes) inside, *b* is a 2*k*-bit offset, and *a* is an odd multiplier. Picking a “random” hash function means picking random values for *a* and *b*.

##### Lemma 1.

*For any* 2*k-bit number b and any odd* 2*k-bit number a, the function g*_*a,b*_ *is a bijection on K* := {0 .. (4^*k*^ − 1)*}.*

*Proof.* Let *y* = *g*_*a,b*_(*x*). By definition, the range of *g*_*a,b*_ on *K* is a subset of *K*. Because |*K*| is a power of 2 and *a* is odd, the greatest common divisor of |*K*| and *a* is 1, and so there exists a unique multiplicative inverse *a′* of *a* modulo 4^*k*^ =|*K*|, such that *aa′* = 1 (mod 4^*k*^). The other operations (xor *b*, rot_*k*_) are inverses of themselves; so we recover *x* = rot_*k*_([(*a′* · *y*) mod 4^*k*^] xor *b*).

In summary, each stored canonical *k*-mer needs 2 +3 + ⌈2*k* − log_2_ − *p*⌉ bits to remember the hash function choice and to store the value (species), and the quotient, respectively. For *k* = 25 and *p* = 1276595745 buckets, this amounts to 25 bits per *k*-mer, or 100 bits for each bucket of 4 *k*-mers. To ensure cache line aligned pages, we could insert 28 padding bits to grow the bucket size to 128 bits; however, we chose less memory for a small speed decrease, and let some buckets cross cache line boundaries.

### 2.4 Annotating weak *k*-mers

A *k*-mer that occurs only in the host (graft) reference, but has a Hamming-distance-1 neighbor in the graft (host) reference, is called a *weak* host (graft) *k*-mer. So for a weak *k*-mer, a single nucleotide variation could flip its assigned species, while a *k*-mer that is not weak is more robust in this sense. A similar concept exists in *xenome*; however, weak host and graft *k*-mers are combined into “marginal” *k*-mers, and their origin is not stored. After the hash table has been constructed with all *k*-mers and their values “host”, “graft” or “both”, we mark weak *k*-mers by modifying the value, setting an additional “weak” bit. In principle, we could scan over the *k*-mers and query all 3*k* neighbors of each *k*-mer, but this is inefficient.

Instead, we extract from the hash table a complete list *L* of *k*-mers and their reverse complements (not canonical codes; approx. 9 10^9^ entries for 4.5 10^9^ distinct *k*-mers), together with their current values. To save memory, this list is created and processed in 16 chunks according to the first two nucleotides of the *k*-mer, thus needing approx. 4.5 GB of additional memory temporarily. Since we use odd *k* = 2*ℓ* + 1, we can partition a *k*-mer into its *ℓ*-prefix, its middle base and its *ℓ*-suffix. We make use of the following observation.

#### Observation 1.

*For k* = 2*ℓ* + 1, *two k-mers x, y with Hamming distance 1 differ either in their ℓ-suffix, in the ℓ-suffix of their reverse complement or in their middle base.*

We thus partition the sorted list into blocks of constant (*ℓ* + 1)-prefixes. Different blocks are processed independently in parallel threads. The *ℓ*-suffixes of all pairs of *k*-mers in such a block are queried with a fast bit-vector test for Hamming distance 1. If a pair is found and the *k*-mers occur in different species, the “weak bit” (value 4) is set. It remains to find pairs of *k*-mers that differ only in their middle base. We conceptually partition the list into blocks of constant *ℓ*-prefixes and use that such pairs must occur consecutively in a block and agree in the *ℓ*-suffix. So these pairs can be identified within a single linear scan. In the end, updated values are transferred to the values of the canonical *k*-mers in the hash table.

### 2.5 Reference sequences

To build the *k*-mer hash table from genomic and transcribed sequences from human and mouse, we obtained the “toplevel DNA” genome FASTA files, which include both the primary assembly, unplaced contigs and alternative alleles, and the “all cDNA” files, which contain the known transcripts, from the ensembl FTP site, release 98.

As the alternative alleles of the human and mouse toplevel references contain mostly Ns to keep positional alignment of alternative alleles to the consensus reference, they decompress to huge FASTA files (over 60 GB for human, over 12 GB for mouse). Therefore we condensed the toplevel reference sequences by replacing runs of more than 25 Ns by 25 Ns. This does not change the *k*-mer content, as *k*-mers containing even a single N are ignored. It does provide an efficiency boost to alignment-based tools because read mappers build an index of every position in the genome and typically replace runs of Ns by random sequence.

We used the following files with the following compressed file sizes and number of contained basepairs, with numbers after ↦ indicating sizes after condensing Ns:

- Graft (human) genome (1100 MB; 61 Gbp ↦ 885 MB; 3.15 Gbp): ftp://ftp.ensembl.org/pub/release-98/fasta/homo_sapiens/dna/Homo_sapiens.GRCh38.dna.toplevel.fa.gz
- Graft (human) transcriptome (67 MB; 0.37 Gbp): ftp://ftp.ensembl.org/pub/release-98/fasta/homo_sapiens/cdna/Homo_sapiens.GRCh38.cdna.all.fa.gz
- Host (mouse) genome (804 MB; 12 Gbp ↦ 776 MB; 2.72 Gbp): ftp://ftp.ensembl.org/pub/release-98/fasta/mus_musculus/dna/Mus_musculus.GRCm38.dna.toplevel.fa.gz
- Host (mouse) transcriptome (50 MB; 0.25 Gbp): ftp://ftp.ensembl.org/pub/release-98/fasta/mus_musculus/cdna/Mus_musculus.GRCm38.cdna.all.fa.gz

### 2.6 Fragment classification

Given a sequenced fragment (single read or read pair), we query each *k*-mer of the fragment about its origins; *k*-mers with undetermined bases are ignored. Our implementation reads large chunks (several MB) of FASTQ files and distributes read classification over several threads (we found that 8 threads saturate the I/O).

We collect *k*-mer statistics for each fragment (adding the numbers of both reads for a read pair): Let *n* be the number of (valid) *k*-mers in the fragment. Let *h* be the number of host *k*-mers and *h′* the number of weak host *k*-mers, and analogously define *g* and *g′* for the graft species. Further, let *b* be the number of *k*-mers occuring in both species, and let *x* be the number of *k*-mers that were not found in the key-value store. Based on the vector (*h, h′, g, g′, b, x*; *n*), we use a tree of hierarchical rules to classify the fragment into one of five categories: “host”, “graft”, “both”, “neither” and “ambiguous”. Categories “host” and “graft” are for reads that can be clearly assigned to one of the species. Category “both” is for reads that match equally well to both references. Category “neither” is for reads that contain many *k*-mers that cannot be found in the key-value store; these could point to technical problems (primer dimers) or contamination of the sample with other species. Finally, category “ambiguous” is for reads that provide conflicting information. Such reads should not usually be seen; they could result from PCR hybrids between host and graft during library preparation. The precise rules are shown in Figure 2. Category “ambiguous” is chosen if no “else” rule exists and no other rule applies in any given node.

**Fig. 2.**
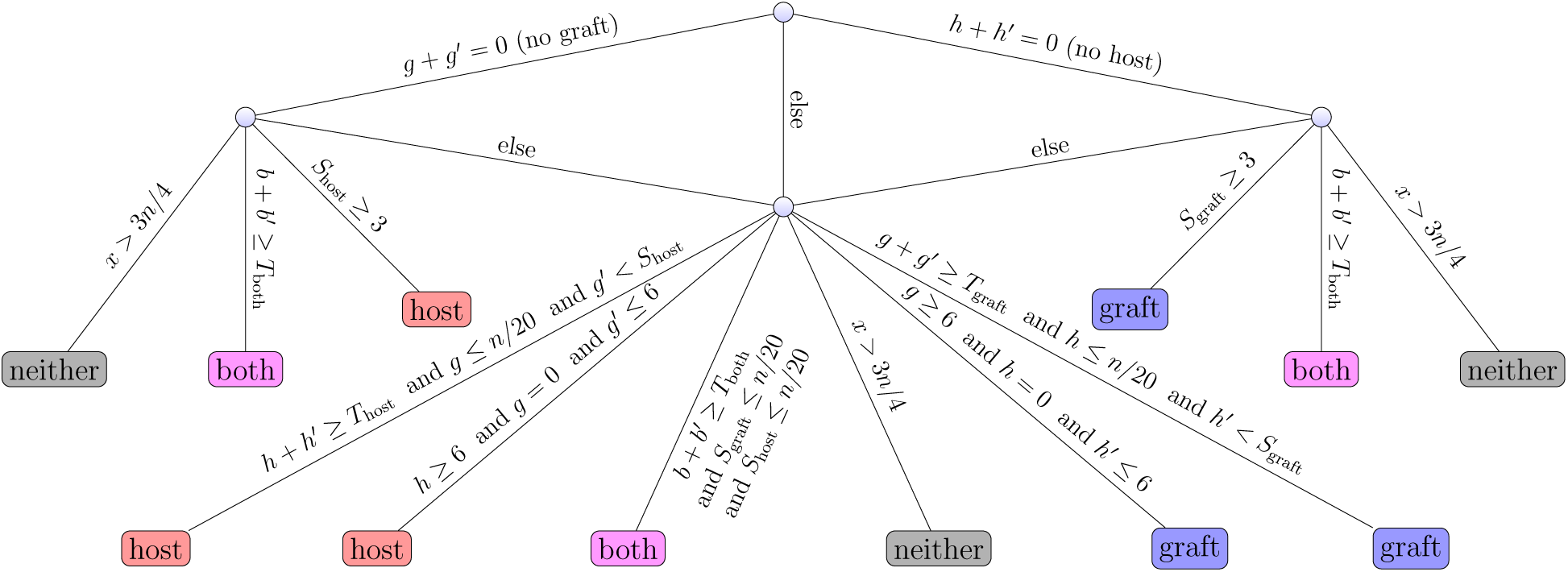
Decision rule tree for classifying a DNA fragment from *k*-mer statistics (*h, h′, g, g′, b, x*; *n*), meaning number of *k*-mers of type “host” (*h*), “weak host” (*h*), “graft” (*g*), “weak graft” (*g*), “both” (*b*), and number of *k*-mers not present in the key-value store (*x*), respectively; *n* is the total number of (valid) *k*-mers in the fragment. We also use weighted scores *S*_host_ := *h* + ⌊*h′/*2⌋ and *S*_graft_ := *g* + ⌊*g′/*2⌋ and thresholds *T*_host_ := ⌊*n/*4⌋, *T*_graft_ := ⌊*n/*4⌋ and *T*_both_ := ⌊*n/*4⌋. A fragment is thus classified as “host”, “graft”, “both”, “neither”, or “ambiguous”. Category “ambiguous” is chosen if no other rule applies and no “else” rule is present in a node.

#### Quick mode

Inspired by a similar acceleration in the *kallisto* software (12) for transcript expression quantification, we additionally implemented a “quick mode” that initially looks only at the type of the third and third-last *k*-mer in every read. If the two (for single-end reads) or four (for paired-end reads) types agree (e.g. all are “graft”), the fragment is classified on this sampled evidence alone. This results in quicker processing of large FASTQ files, but only considers a small sample of the available information.

## 3 Results

We evaluate our alignment-free xenograft sorting approach and its implementation *xengsort* for the common case of human-tumor-in-mouse xenografts, by using mouse datasets, human datasets, xenograft datasets and datasets from other species, and compare against an existing tool with the same purpose, *xenome* from the *gossamer* suite (7), and against a representative of alignment-based filtering tools, *XenofilteR* (2). The hardware used for the benchmarks was one server with two AMD Epyc 7452 CPUs (with 32 cores and 64 threads each), 1024 GB DDR4-2666 memory and one 12 TB HDD with 7200 rpm and 256 MB cache.

### 3.1 Hash table construction

#### Table size and uniqueness of *k*-mers

We evaluated *k* ∈ {23, 25, 27} and then decided to use *k* = 25 because it offers a good compromise between species specificity and memory requirements. Table 2 shows several index properties. In particular, moving from *k* = 25 to *k* = 27, the small decrease in *k*-mers that map to both genomes and in weak *k*-mers did not justify the additional memory requirements. In addition, longer *k*-mers lead to lower error tolerance against sequencing errors, as each error affects up to *k* of the *k*-mers in a read.

**Table 2.**
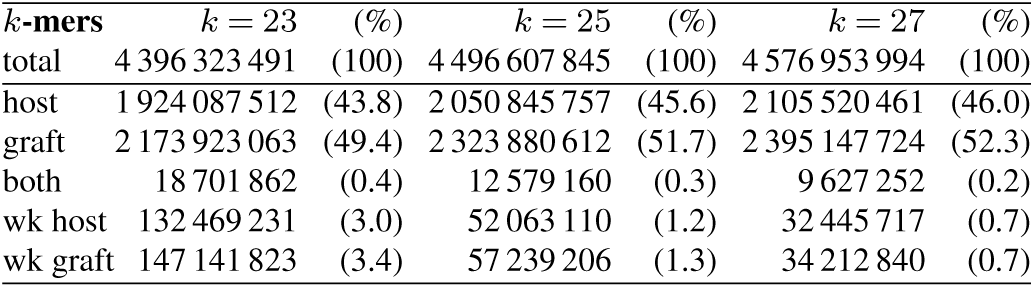
Properties of the *k*-mer index for different values of *k* (wk: weak). Underlying reference sequences are given in Section 2.5.

#### Construction time and memory

Table 3 shows time and memory requirements for building the *k*-mer hash table or FM index for *bwa* (for *XenofilteR*). The main difference is that the BWA index is a succinct representation of the suffix array of the references and not a *k*-mer hash table. Our hash table construction is not paralellized; hence CPU times and wall clock times agree and are less than one hour. The hash construction of *xenome* is paralellized; we gave it 8 threads (but 9 were sometimes used); yet it does about 20 times the CPU work and takes three times as long as *xengsort*, even when using multiple threads.

**Table 3.**
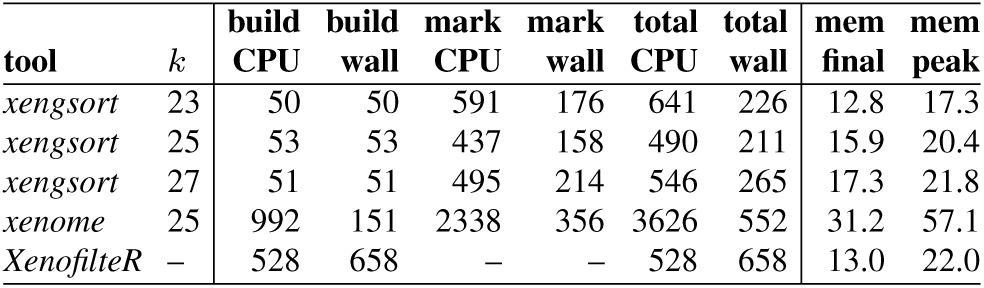
Index construction: CPU times and wall clock times in minutes and memory in Gigabytes using different tools and different *k*-mer sizes for *xengsort*. “Build” times refer to collecting and hashing the *k*-mers according to species, but without marking weak *k*-mers. “Mark” times refer to marking weak *k*-mers. “Total” times are the sum of build and mark times, plus additional I/O times. “CPU” times measure total CPU work load (as reported by the time command as user time), and “wall” times refer to actually passed time. Final size (“mem final”) is measured by index size on disk (GB). Memory peak (“mem peak”) is the highest memory usage during construction (GB).

Marking weak or marginal *k*-mers is paralellized in both approaches; wall clock times are measured using 8 threads.

Again, *xengsort* finds the weak *k*-mers faster, both in terms of total CPU work and wall clock time.

The indexing method of bwa is not comparable, as it builds a complete suffix array (FM index) that is independent of *k* and does not include marking weak *k*-mers. Here the CPU time is lower than the wall clock time, which indicates an I/O starved process.

We note that *xenome* uses a large amount of memory during hash table construction (it was given up to 64 GB). It works with less if restricted, but at the expense of longer running times. BWA indexing also needs significant additional memory during construction. The additional memory required by *xengsort* results from the additional sorted *k*-mer list required for detecting weak *k*-mers. Overall, our construction is fast (even though serial only) and uses a reasonable amount of memory.

#### Load factor and hash choice distribution

As explained in Section 2.3, 3-way Cuckoo hash tables support very high loads (fill ratios) over 99.9%. However, such loads come at the expense of distributing all *k*-mers almost evenly across hash function choices. For faster lookup, it is beneficial to leave part of the hash table empty. We used a load factor of 88% and thus find 76.7% of the *k*-mers at their first bucket choice, 15.5% at their second choice and only 7.8% at their third choice, yielding an average of 1.31 lookups for a present *k*-mer.

Applying assignment optimization (11), which takes an additional 5 hours (serial CPU time, not parallelized) and needs over 80 GB of RAM, we achieve a slightly better average of 1.17 lookups for a present *k*-mer.

### 3.2 Classification results

We applied our method *xengsort, xenome* and *XenofilteR* to several datasets with reads of known origin (except possible contamination issues or technical artefacts), that however present certain particular challenges. A summary of running times for all datasets appears in Table 4.

**Table 4.** Dataset sizes (number of fragments; M: millions) and CPU times in minutes spent on different datasets, measured with the “time” command (user time) when running with 8 threads (*xenome, xengsort, bwa-mem*, BAM sorting, except for *XenofilteR* (XfR), which is single-threaded). N/A: not applicable; tool could not be run on this dataset.

| dataset / tool | size | XfR + | bwa + | sort | *xenome* | *xengsort* |
| --- | --- | --- | --- | --- | --- | --- |
| mouse exomes | 307 M | 310 + | 8291 + | 179 | 1823 | 368 |
| human matepair | 1258 M | N/A + | 222939 + | 940 | 9845 | 2463 |
| chicken genome | 251 M | 76 + | 6976 + | 118 | 1273 | 592 |
| leukemia RNA | 1760 M | 778 + | 22111 + | 521 | 5188 | 1680 |
| PDX RNA | 9742 M | 16043 + | 278329 + | 5862 | 59692 | 13555 |

#### Human-captured mouse exomes

A recent comparative study (1) made five mouse exomes accessible, which were captured with a human-exome capture kit and hence presents mouse reads that are biased towards high similarity with human reads. The mouse strains were A/J (two mice), BALB/c (one mouse), and C57BL6 (two mice); they were sequenced on the Illumina HiSeq 2500 platform, resulting in 11.8 to 12.7 Gbp. The datasets are available under accession numbers SRX5904321 (strain A/J, mouse 1), SRX5904320 (strain A/J, mouse 2), SRX5904319 (strain BALB/c, mouse 1), SRX5904318 (strain C57BL/6, mouse 1) and SRX5904322 (strain C57BL/6, mouse 2).

Ideally, all reads should be classified as mouse reads. Table 5 shows detailed classification results and running times. Considering the BALB/c and C57BL/6 strains first, it is evident that classification accuracy is high (over 98.9% mouse for *xengsort*, over 97.4% for *xenome*; with less than 0.64% human reads for both tools). The main difference between the tools is that *xenome* is more conservative, assigning a larger fraction of reads to the “ambiguous” (unclassified) category. With *xenome*, this happens for reads that contain two *k*-mers *x, y*, where *x* maps uniquely to human and *y* maps uniquely to mouse. The decision rule of *xengsort* is more permissive and tolerant towards small inconsistencies. Therefore, *xengsort* assigns more reads correctly to mouse, and fewer to the ambiguous category. Additionally, *xengsort* assigns fewer reads incorrectly to human.

**Table 5.** Detailed classification results on five human-captured mouse exomes from different mouse strains (2*×* A/J, 1*×* BALB/c, 2*×* C57BL/6). Running times are reported both in CPU minutes [Cm], measuring CPU work, and wall clock minutes [Wm], measuring actual time spent. Times for *XenofilteR* (XfR) do not include alignment or BAM sorting time. Classification results report the number and percentage (in brackets) of fragments classified as mouse (correct), both human and mouse (likely correct), human (incorrect), ambiguous (no statement) and neither (likely incorrect). *XenofilteR* (XfR) only extracts human fragments and does not classify the remainder; so only the number of fragments classified as human are reported.

| A/J-1 | *xengsort* |  | *xenome* |  | XfR |  |
| --- | --- | --- | --- | --- | --- | --- |
|  | 70 Cm | 14 Wm | 371 Cm | 45 Wm | 56 Cm | 56 Wm |
|  | fragments | (%) | fragments | (%) | fragments | (%) |
| mouse | 46 648 014 | (78.03) | 45 759 814 | (76.54) |  |  |
| both | 120 808 | (0.20) | 65 269 | (0.11) |  |  |
| human | 12 813 583 | (21.43) | 12 500 844 | (20.91) | 6 315 955 | (10.56) |
| ambgs. | 58 449 | (0.10) | 1 383 547 | (2.31) |  |  |
| neither | 143 775 | (0.24) | 75 155 | (0.13) |  |  |

| A/J-2 | *xengsort* |  | *xenome* |  | XfR |  |
| --- | --- | --- | --- | --- | --- | --- |
|  | 70 Cm | 15 Wm | 416 Cm | 50 Wm | 67 Cm | 67 Cm |
| mouse | 60 255 189 | (95.57) | 59 135 489 | (93.80) |  |  |
| both | 151 396 | (0.24) | 89 089 | (0.14) |  |  |
| human | 2 301 384 | (3.65) | 2 271 131 | (3.60) | 1 718 545 | (2.73) |
| ambgs. | 57 827 | (0.09) | 1 340 814 | (2.13) |  |  |
| neither | 279 556 | (0.44) | 208 829 | (0.33) |  |  |

| BALB/c | *xengsort* |  | *xenome* |  | XfR |  |
| --- | --- | --- | --- | --- | --- | --- |
|  | 68 Cm | 15 Wm | 392 Cm | 45 Wm | 61 Cm | 61 Wm |
| mouse | 62 235 960 | (98.99) | 61 274 277 | (97.46) |  |  |
| both | 118 541 | (0.19) | 68 949 | (0.11) |  |  |
| human | 342 908 | (0.55) | 348 154 | (0.55) | 285 556 | (0.45) |
| ambgs. | 45 063 | (0.07) | 1 098 036 | (1.65) |  |  |
| neither | 127 035 | (0.20) | 80 091 | (0.13) |  |  |

| C57BL/6-1 | *xengsort* |  | *xenome* |  | XfR |  |
| --- | --- | --- | --- | --- | --- | --- |
|  | 72 Wm | 14 Wm | 359 Wm | 44 Wm | 58 Cm | 58 Wm |
| mouse | 57 993 361 | (98.93) | 57 522 446 | (98.13) |  |  |
| both | 118 984 | (0.20) | 74 325 | (0.13) |  |  |
| human | 375 716 | (0.64) | 376 653 | (0.64) | 290 894 | (0.50) |
| ambgs. | 27 731 | (0.05) | 571 542 | (0.98) |  |  |
| neither | 103 895 | (0.18) | 74 721 | (0.13) |  |  |

| C57BL/6-2 | *xengsort* |  | *xenome* |  | XfR |  |
| --- | --- | --- | --- | --- | --- | --- |
|  | 67 Cm | 15 Wm | 422 Cm | 51 Wm | 62 Cm | 62 Wm |
| mouse | 62 384 448 | (99.00) | 61 941 783 | (98.30) |  |  |
| both | 107 019 | (0.17) | 66 163 | (0.10) |  |  |
| human | 189 536 | (0.30) | 208 149 | (0.33) | 132 535 | (0.21) |
| ambgs. | 27 142 | (0.04) | 562 659 | (0.89) |  |  |
| neither | 304 677 | (0.48) | 234 068 | (0.37) |  |  |

However, the two samples of strain A/J give different results. Both *xengsort* and *xenome* assign a large fraction of reads (around 21% and 3.6% in the two samples) to the human genome, while *XenofilteR* assigns only 10.5% and 2.7%, respectively. While *xengsort* does assign more reads to mouse, it also assigns more reads to human, following its strategy of leaving fewer reads unassigned (ambiguous). Inspection of these reads revealed that almost all of them are low-complexity, i.e. consist of repetitive sequence, and a check with BLAT (13) revealed no hits in mouse and several gapped hits in the human genome. So the classification as human reads is not incorrect from a technical standpoint, but in fact these reads appear to point to techincal problems during then enrichment step of the library generation. An additional low-complexity filter would remove most problematic reads. Concerning running times, we find that *xengsort* needs around 70 CPU minutes for one of these datasets, and less than 15 minutes of wall clock time using 8 threads. The speed-up being less than 8 results from serial intermediate I/O steps. While *xenome* makes better use of parallelism, it is slower overall, requiring 5 to 6 times the CPU work of *xenome*. For only scanning already aligned BAM files, *XenofilteR* is surprisingly slow, and we see that we can sort the reads from scratch in almost the same amount of CPU work that is required to compare alignment scores. When adding bwa mem alignment times (even without the time required for sorting the resulting BAM files), *XenofilteR* needs an additional 887 to 1424 CPU minutes for the human alignments and an additional 424 to 777 minutes for the mouse alignments per dataset, making the alignment-based approach far less efficient than the alignment-free approach.

#### Human genome (GIAB) matepair library

We obtained FASTQ files of an Illumina-sequenced 6kb matepair library from the Genome In A Bottle (GIAB) Ashkenazim trio dataset according to the provided sequence file index (ftp://ftp-trace.ncbi.nlm.nih.gov/giab/ftp/data_indexes/AshkenazimTrio/sequence.index.AJtrio_Illumina_6kb_matepair_wgs_08032015). The data represents a family (mother, father, son). Ideally, we see only human reads.

Figure 3A shows the classification results for *xengsort* and *xenome. XenofilteR* reported that the BAM files were too large to be processed and did not give a result (400 GB total for human and mouse; each BAM file over 30 GB in size). We see that almost all reads are correctly identified as human, while a small fraction is neither, which could be adapter dimers or other technical issues. However, *xenome* classifies a similarly small fraction as ambiguous. We observe the same wall clock time ratio (about 3.5) between *xenome* and *xengsort* as for the mouse exome dataset.

**Fig. 3.**
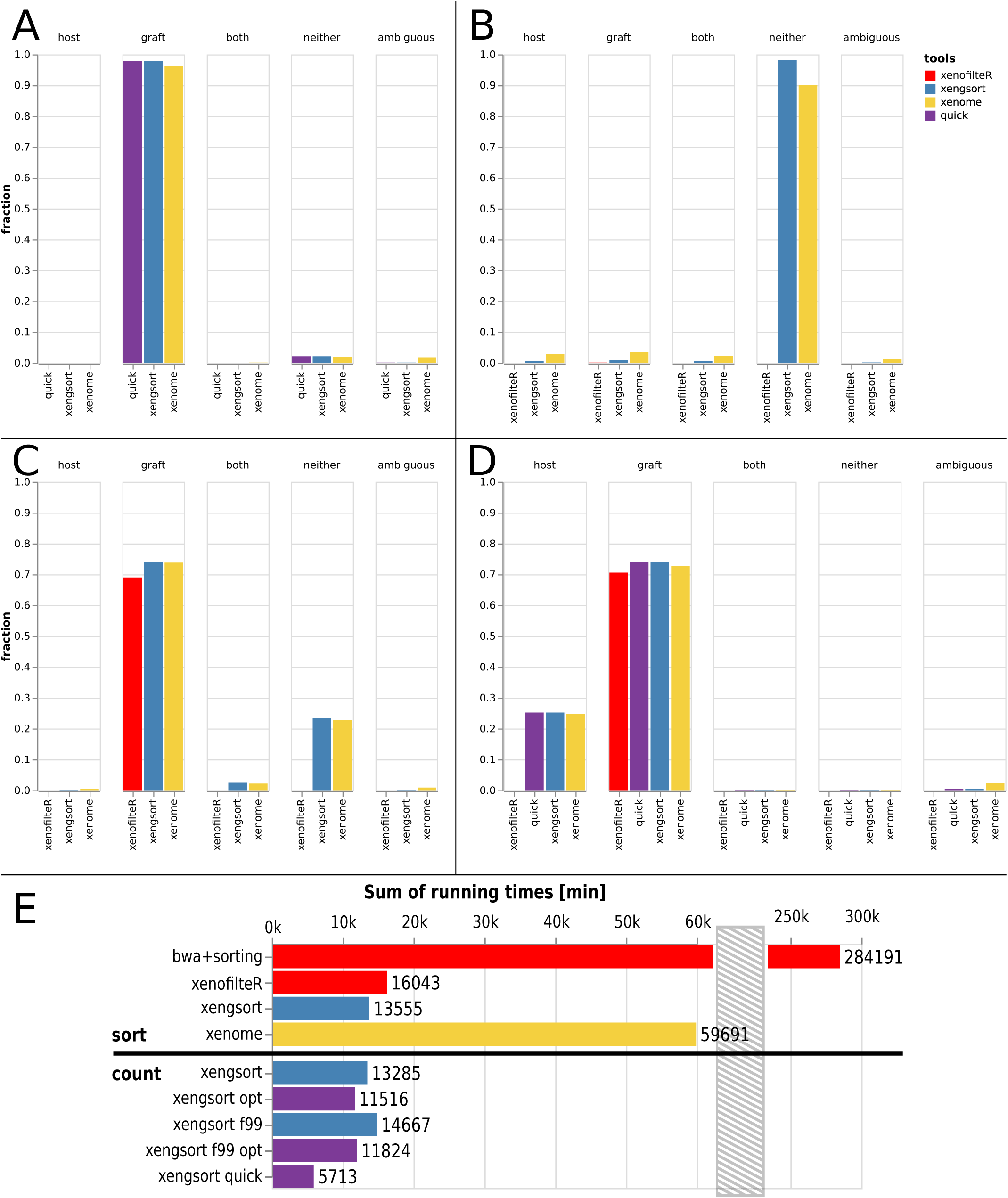
Classification results of different tools (*XenofilteR, xenome, xengsort*, and partially *xengsort* with “quick” option) on several datasets: A: GIAB human matepair dataset (*XenofilteR* did not run on this dataset); B: Chicken genome; C: Human lymphocytic leukemia RNA-seq data; D: Patient-derived xenograft (PDX) RNA-seq data. E: CPU times on the PDX RNA-seq dataset with different tools and different *xengsort* parameters (see text).

Because this is a very large dataset (112 GB gzipped FASTQ), we additionally evaluated the effects of using *xengsort*’s “quick mode”. We observed a significant reduction in processing time (by about 33%) and almost unchanged classification results. We also ran the *xengsort* classification with the optimized hash table (using an optimized assignment computed using the methods from (11) and found a small reduction (9%) in running time.

We conclude that both alignment-free tools accurately recognize that this is a pure human dataset, and that *xengsort* is again more CPU-efficient and faster, given the same resources.

#### Chicken genome

We obtained a paired-end (2×101bp) Illumina whole genome sequencing run of a chicken genome from a whole blood sample (accession SRX6911418) with a total of 251 million paired-end reads. Ideally, none of these reads are recognized as mouse or human reads. Figure 3B shows divergent results. For *XenofilteR*, we can only say that almost no reads are extracted as human; the remainder is unclassified. *Xenome* assigns a small number of reads to each category and only around 90% into the “neither” category, while *xengsort* assigns 98.11% of the reads as “neither”.

Concerning running time (cf. Table 4), the scan of *XenofilteR* here beats the alignment-free tools because both BAM files are essentially empty, as very few reads align against human or mouse. Also, the speed advantage of *xengsort* over *xenome* is less on this dataset, mainly because most *k*-mers are not found in the index and require *h* = 3 memory lookups and likely cache misses. Such a dataset that contains neither graft nor host reads is aversarial for our design of *xengsort*; it is also unlikely to be encountered in practice.

#### Human lymphocytic leukemia tumor RNA-seq data

We obtained single-end FASTQ files from RNA-seq data of 5 human T-cell large granular lymphocytic leukemia samples, where recurrent alterations of TNFAIP3 were observed, and 5 matched controls (13.4 Gbp to 27.5 Gbp). The files are available from SRA accession SRP059322 (datasets SRX1055051 to SRX1055060). Surprisingly, not all fragments were recognized as originating from human tissue (Figure 3C). While *xenome* and *xengsort* agreed that the human fraction is close to 75%, *XenofilteR* assigned considerably fewer reads to human origins (less than 70%).

For this and other RNA-seq datasets, we trimmed the Illumina adapters using cutadapt (14) prior to classification, as some RNA fragments may be shorter than the read length. If this step is omitted, even fewer fragments are classified as human (graft): just below 70% for *xenome* and *xengsort*, and only about 53% for *XenofilteR*. The number of fragments classified as neither increases correspondingly.

We investigated the reads classified by *xengsort* as neither human nor mouse. Quality control with FastQC (15) revealed nothing of concern, but showed an unusual biomodal per-fragment GC content distribution with peaks at 45% and 55%. BLASTing the fragments against the non-redundant nucleotide database (16) yielded no hits at all for 97% of these fragments. A small number (2%) originated from the bacteriophage PhiX, which was to be expected, because it is a typical spike-in for Illumina libraries. The remaining 1% of fragments showed random hits over many species without a distinctive pattern. We therefore concluded that the neither fragments mainly consisted of artefacts from library construction, such as ligated and then sequenced random primers.

Concerning running times (Table 4), we observed again that *xengsort* is more than 3 times faster than *xenome*and that *xengsort* needs time comparable to *XenofilteR*even when only the time for sorting and scanning existing BAM files is taken into account. Producing the alignments takes much longer.

#### Patient-derived xenograft (PDX) RNA-seq samples from human pancreatic tumors

We evaluated 174 pancreatic tumor patient-derived xenograft (PDX) RNA-seq samples that are available internally at University Hospital Essen. Figure 3D shows that all three tools classify between 70% and 74% as graft (human) fragments. Again, *XenofilteR* seems to be the most conservative tool with about 70%, and *xenome* classifies about 72% as human and *xengsort* 74%. The remaining reads are not classified by *XenofilteR*, while *xenome* and *xengsort* both assign about 25% to host (mouse). Furthermore, *xenome* classifies about 2% and *xengsort* less than 1% as ambiguous. So we observe that on all datasets, *xengsort* is more decisive than *xenome* and, judging from the pure human and mouse datasets, mostly correct about it. Because this is a large dataset, we also applied *xengsort*’s quick mode and found essentially no differences in classification results (less than 0.001 percentage points in each class; e.g. for graft: quick 74.0111% vs. standard 74.0105% of all reads; difference 0.0006%; cf. Fig. 3D).

Concerning running time, Figure 3E shows that the alignment using *bwa-mem* and the sorting of the BAM file for *XenofilteR* took over 284 191 CPU minutes (close to 200 days). After that, *XenofilteR* required an additional 16 043 CPU minutes (over 11 days) to classify the aligned and sorted reads. In comparison, *xenome* with 59 691 CPU minutes (41.5 days) took only 20% of the time used by *bwa-mem* and *XenofilteR*, and *xengsort* needed 13 555 CPU minutes (9.5 CPU days) to sort all reads and is therefore even faster than the classification by *XenofilteR* alone, even excluding the alignment and sorting steps, and over 4 times faster than *xenome*. Using the “quick mode” with an optimized hash table at 88% load needed only 5 713 CPU minutes (less than 4 CPU days), i.e., less than half of the time of a full analysis.

We additionally examined some trade-offs for this dataset. First, we note that only counting proportions without output (“count” operation) is not much faster than sorting the reads into different output files (“sort” operation): 13285 vs. 13555 CPU minutes (2% faster). We additionally measured the running time of the *xengsort*’s count operation on hash tables with different load factors (88% and 99%) using both the standard assignment by random walk and an optimal assignment (11). As expected, a load factor of 99% was slower than 88% (by 10.4% on the random walk assignment, but only by 2.6% on the optimized assignment). Using the optimal assignment gives a speed boost (13.3% faster at 88% load; 19.3% at 99% load). The optimized assignment at 99% load yields an even faster running time than the random walk assignment at 88% load by 11% (11 824 vs. 13 285 CPU minutes).

## 4 Discussion and Conclusion

We revisited the xenograft sorting problem and improved upon the state of the art in alignment-free methods with our implementation of *xengsort*.

On typical datasets (PDX RNA-seq), it is is at least four times faster and needs less memory than the comparable *xenome* tool. Our experiments show that it provides accurate classification results, and classifies more reads than *xenome*, which more often bails out when uncertain. Surprisingly, on PDX datasets, our approach is even faster than scanning already aligned BAM files. This favorable behavior arises because almost every *k*-mer in every read can be expected to be found in the key-value store, and lookups of present keys are faster than lookups of absent keys with our data structure.

On adversarial datasets (e.g., a sequenced chicken genome, where almost none of the *k*-mers can be found in the hash table), *xengsort* is 2 times faster than *xenome* and about 8 times slower than scanning pre-aligned and pre-sorted BAM files (which are mostly empty).

However, given that producing and sorting the BAM files takes significant additional time, especially for computing the (non-existing) alignments, our results show that overall, alignment-free methods require significantly less computational resources than alignment-based methods. In view of the current worldwide discussions on climate change and energy efficiency, we advocate that the most resource-efficient available methods should be used for a task, and we propose that *xengsort* is preferable to existing work in this regard. Even though one could argue that alignments are needed later anyway, we find that this is not always true: First, to analyze PDX samples, typically only the graft reads are further considered and need to be aligned. Second, recent research has shown that more and more application areas can be addressed by alignment-free methods, even structural variation and variant calling (17), so alignments may not be needed at all.

On the methodological side, we developed a general key-value store for DNA/RNA *k*-mers that allows extremely fast lookups, often only a single random memory access, and that has a low memory footprint thanks to a high load factor and the technique of quotienting.

Thus this work might be seen as a blueprint for implementations of other alignment-free methods (for gene expression quantification, metagenomics, etc.). In principle, one could replace the underlying key-value store of each published *k*-mer based method by the hashing approach presented here and probably obtain a speed-up of factor 2 to 4, while at the same time saving some space for the hash table. In practice, such an approach may be difficult because the code in question is often deeply nested in the application. However, we would like to suggest that for future implementations, three-way bucketed Cuckoo hash tables with quotienting should be given serious consideration.

A (small) limitation of our approach is that the size of the hash table must be known (at least approximately) in advance. (Growing it would mean re-hashing everything). Fortunately, the total length of the sequences in the *k*-mer key-value store provides an easily calculated upper bound. The advantage of such a static approach is that only little additional memory is required during construction.

The software *xengsort* is available at http://gitlab.com/genomeinformatics/xengsort under the MIT license. Installation and usage instructions are provided within the README file of the repository. The software is written in Python, but makes use of just-in-time compilation at runtime using the numba package (18). While requiring an additional 1–2 seconds of startup time, this allows for many optimizations, because certain parameters that become only known at run time, such as random parameters for the hash functions, can be compiled as constants into the code. These optimizations yield savings that can exceed the initial compilation effort.

Further variants of our approach can be explored and evaluated: We already introduced a “quick mode”, similar to the one in kallisto (12), that is faster, but may falsely classify problematic (amiguous) reads as belonging to a specific species. In practice, this does not appear to be a problem. In the future, we may alternatively reduce the number of *k*-mer lookups by not examining every *k*-mer, but only minimizers in windows of fixed size, using min-hashing or other sampling methods. Another alternative is to base the classification not on the number of (overlapping) *k*-mers belonging to each species, but on the number of basepairs covered by *k*-mers of each species. Such investigations are ongoing.

While we have indications that classification results agree well overall among all methods and variants, we concur with a recent study (1) that there exist subtle differences, whose effects can propagate through computational pipelines and influence, for example, variant calling results downstream, and we believe that further evaluation studies are necessary. In contrast to their study, we however suggest that a best practice workflow for PDX analysis should start (after quality control and adapter trimming on RNA-seq data) with *alignment-free* xenograft sorting, followed by aligning the graft reads and the reads that can originate from both genomes to the graft genome. In any workflow, the latter reads, classified as “both”, may pose problems, because one may not be able to decide the species of origin. Indeed, ultraconserved regions of DNA sequence exist between human and mouse. In this sense we believe that full read sorting (into categories host, graft, both, neither, ambiguous, as opposed to extracting graft reads only) gives the highest flexibility for downstream steps and is prefereable to filter-only apporaches.

## Acknowledgements

This work was funded by the Mercator Research Center Ruhr (MERCUR), project Pe-2013-0012 (UA Ruhr professorship) and by the German Research Foundation (DFG), Collaborative Research Center SFB 876, project C1.

## References

1. S. Y. Jo, E. Kim, and S. Kim. Impact of mouse contamination in genomic profiling of patient-derived models and best practice for robust analysis. Genome Biology, 20(1):Article 231, Nov 2019.

2. R. J. C. Kluin, K. Kemper, T. Kuilman, J. R. de Ruiter, V. Iyer, J. V. Forment, P. Cornelissen-Steijger, I. de Rink, P. Ter Brugge, J. Y. Song, S. Klarenbeek, U. McDermott, J. Jonkers, A. Velds, D. J. Adams, D. S. Peeper, and O. Krijgsman. XenofilteR: computational deconvolution of mouse and human reads in tumor xenograft sequence data. BMC Bioinformatics, 19(1):366, Oct 2018.

3. Gnöknur Giner. XenoSplit, 2019. Unpublished; source code available at https://github.com/goknurginer/XenoSplit.

4. G. Khandelwal, M. R. Girotti, C. Smowton, S. Taylor, C. Wirth, M. Dynowski, K. K. Frese, G. Brady, C. Dive, R. Marais, and C. Miller. Next-generation sequencing analysis and algorithms for PDX and CDX models. Mol. Cancer Res., 15(8):1012–1016, 08 2017.

5. M. J. Ahdesmäki, S. R. Gray, J. H. Johnson, and Z. Lai. Disambiguate: An open-source application for disambiguating two species in next generation sequencing data from grafted samples. F1000Res, 5:2741, 2016.

6. Brian Bushnell. BBsplit, 2014–2020. Part of BBTools, https://jgi.doe.gov/data-and-tools/bbtools/.

7. T. Conway, J. Wazny, A. Bromage, M. Tymms, D. Sooraj, E. D. Williams, and B. Beresford-Smith. Xenome–a tool for classifying reads from xenograft samples. Bioinformatics, 28(12):i172–i178, Jun 2012.

8. M. Callari, A. S. Batra, R. N. Batra, S. J. Sammut, W. Greenwood, H. Clifford, C. Hercus, S. F. Chin, A. Bruna, O. M. Rueda, and C. Caldas. Computational approach to discriminate human and mouse sequences in patient-derived tumour xenografts. BMC Genomics, 19(1):19, 2018.

9. W. Dai, J. Liu, Q. Li, W. Liu, Y. X. Li, and Y. Y. Li. A comparison of next-generation sequencing analysis methods for cancer xenograft samples. J Genet Genomics, 45(7):345–350, 2018.

10. Stefan Walzer. Load thresholds for cuckoo hashing with overlapping blocks. In Ioannis Chatzigiannakis, Christos Kaklamanis, Dániel Marx, and Donald Sannella, editors, 45th International Colloquium on Automata, Languages, and Programming, ICALP 2018, volume 107 of LIPIcs, pages 102:1–102:10. Schloss Dagstuhl - Leibniz-Zentrum fuer Informatik, 2018. doi: 10.4230/LIPIcs.ICALP.2018.102.

11. Jens Zentgraf, Henning Timm, and Sven Rahmann. Cost-optimal assignment of elements in genome-scale multi-way bucketed cuckoo hash tables. In Proceedings of the Symposium on Algorithm Engineering and Experiments (ALENEX) 2020, pages 186–198. SIAM, 2020. doi: 10.1137/1.9781611976007.15.

12. N. L. Bray, H. Pimentel, P. Melsted, and L. Pachter. Near-optimal probabilistic RNA-seq quantification. Nat. Biotechnol., 34(5):525–527, 05 2016. Erratum in Nat. Biotechnol. 34(8):888 (2016).

13. W. J. Kent. BLAT–the BLAST-like alignment tool. Genome Res., 12 (4):656–664, Apr 2002.

14. Marcel Martin. Cutadapt removes adapter sequences from highthroughput sequencing reads. EMBnet.journal, 17(1):10–12, May 2011. doi: http://dx.doi.org/10.14806/ej.17.1.200.

15. Simon Andrews. FastQC: A quality control tool for high throughput sequence data, 2010.

16. C. Camacho, G. Coulouris, V. Avagyan, N. Ma, J. Papadopoulos, K. Bealer, and T. L. Madden. BLAST+: architecture and applications. BMC Bioinformatics, 10:421, Dec 2009.

17. D. S. Standage, C. T. Brown, and F. Hormozdiari. Kevlar: A mapping-free framework for accurate discovery of de novo variants. iScience, 18:28–36, Jul 2019.

18. Siu Kwan Lam, Antoine Pitrou, and Stanley Seibert. Numba: a LLVM-based python JIT compiler. In Hal Finkel, editor, Proceedings of the Second Workshop on the LLVM Compiler Infrastructure in HPC, LLVM 2015, pages 7:1–7:6. ACM, 2015. ISBN 978-1-4503-4005-2. doi: 10.1145/2833157.2833162.

